# Near infrared-light treatment alters mitochondrial homeostasis to induce senescence in breast cancer cells

**DOI:** 10.1101/2023.07.06.547935

**Authors:** I Kalampouka, R R Mould, S W Botchway, A Mackenzie, A V Nunn, E L Thomas, J D Bell

**Affiliations:** Research Centre for Optimal Health, School of Life Sciences, University of Westminster; Research Complex at Harwell & Central Laser Facility, Rutherford Appleton Laboratory, Didcot, Oxford, OX11 0QX, England, United Kingdom; The Guy Foundation – The Guy Foundation, Chedington Court, Beaminster, Dorset DT8 3HY, UK

**Keywords:** senescence, near-infrared, breast cancer, photobiomodulation, mitochondria, ROS, quantum biology

## Abstract

The application of near infrared (NIR)-light to living systems has been suggested as a potential method to enhance tissue repair, decrease inflammation, and possibly mitigate cancer therapy-associated side effects. In this study, we examined the effect of exposing three cell lines: breast cancer (MCF7), non-cancer breast cells (MCF10A), and lung fibroblasts (IMR-90), to 734 nm NIR-light for 20 minutes per day for six days, and measuring changes in cellular senescence. Positive senescent populations were induced using doxorubicin. Flow cytometry was used to assess relative levels of senescence together with mitochondria-related variables. Exposure to NIR-light significantly increased the level of senescence in MCF7 cells (13.5%; P<0.01), with no observable effects on MCF10A or IMR-90 cell lines. NIR-induced senescence was associated with significant changes in mitochondria homeostasis, including raised ROS level (36.0%; P<0.05) and mitochondrial membrane potential (14.9%; P<0.05), with no changes in mitochondrial Ca^2+^. These results suggest that NIR-light exposure can significantly arrest the proliferation of breast cancer cells via inducing senescence, while leaving non-cancerous cell lines unaffected.

## INTRODUCTION

Photobiomodulation (PBM) describes changes in cellular activity and transformation in response to exposure to extrinsic light under controlled conditions (Hamblin, 2017). The current term PBM therapy (or low-level laser therapy – LLLT) is the non-thermal use of light that was first introduced by Mester in 1967 (Mester et al., 1968). PBM therapy, which utilizes red to near infrared (NIR)-light (600 – 1000 nm), appears to promote wound healing, reduce inflammation, manage pain and age-related symptoms (Hamblin, 2017), and is becoming increasingly popular due to its ability to penetrate deeper into human tissue than other wavelengths of light below 600 nm (Ash et al., 2017). PBM is different to photo-dynamic therapy where a similar light energy is used together with photo sensitising drug to induce therapy or cell death. The role of PBM in reducing inflammation is the most widely accepted effect of PBM therapy (Hamblin, 2017), and has been shown to reduce the expression of inflammatory marker genes in cells (Pooam et al., 2021), skin (Choi et al., 2011), and experimental models of age-related macular degeneration (Begum et al., 2013). An emerging application of PBM is to mitigate against the side effects of cancer treatment such as chemotherapy (Robjins et al., 2022).

The mechanism through which PBM may have long lasting effects is not fully understood. The mode of action of PBM involves a cellular chromophore absorbing a photon and exciting an electron from a low-energy to a higher-energy orbit (de Sousa, 2017). Karu (2008) identified cytochrome c oxidase (CCO) in the mitochondrial respiratory chain as a primary chromophore and proposed “retrograde mitochondrial signaling” to explain how short exposure to light can result in long-lasting effects (Karu, 2008). Since then, changes in CCO activity have been observed after PBM (Choi et al., 2021), although alternative mechanisms for these observations have been proposed including the potential modulation of water order around mitochondria and/or cellular light-sensitive ion channels (Sommer et al., 2015; Hamblin, 2017).

Cellular senescence describes cells that have permanently lost their ability to proliferate while staying highly metabolically active (Hayflick and Moorhead, 1961), with mitochondria contributing to the senescent phenotype (Vasileiou et al., 2019). The senescent response to stress can be beneficial to many physiological processes, including development, tissue remodelling, and wound healing (He and Sharpless, 2017). Conversely, the same effect during aging can result in detrimental physiological changes, including increased inflammation, and the onset of many non-inheritable diseases (da Silva et al., 2019; McHugh and Gil, 2018).

Given the parallels between the positive effects of PBM and senescence on physiological processes, we investigated the potential of NIR light to modulate cellular senescence. As senescence can both affect and be affected by changing levels of ROS and redox reprogramming, we hypothesise that NIR-light exposure can increase senescence levels through a bidirectional effect that is dependent on the cell type, mediated by ROS and modulated by mitochondrial function.

## RESULTS

### Doxorubicin induces senescence in cancer and non-cancer cell lines

Premature senescence was induced in MCF7, MCF10A, and IMR-90 cell lines following treatment with 0.25 μM doxorubicin (Dox). Seven days after Dox-treatment, the proportion of senescent cells in the population increased by 53.8% ± 7.5 (P<0.0001) in MCF7, 101.0% ± 9.7 (P<0.0001) in MCF10A and 55.0% ± 8.2 (P=0.0001) in IMR-90 cell populations (Figure 1).

**Figure 1:**
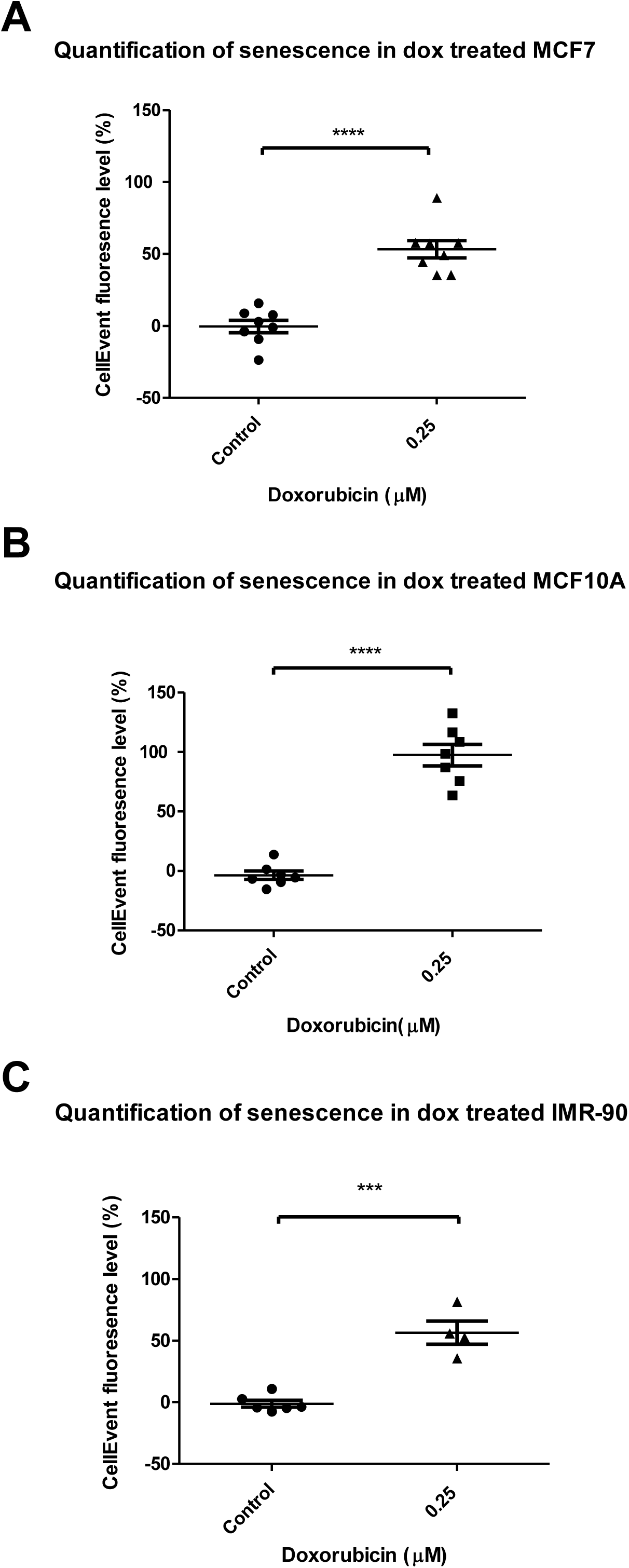
The effect of Dox on cellular senescence induction measured by flow cytometry (A) MCF7 (N=8), (B) MCF10A (N=7) and (C) IMR-90 (N=4) cells were treated with 0.25 μM Dox and the senescent levels were quantified with CellEvent fluorescence intensity. Data are presented as mean changes compared to an untreated control ± SEM. ***: P≤0.001, ****: P ≤0.0001. Control cells followed the same experimental protocol but without the addition of Dox.

### NIR-light treatment increases senescence in both control and Dox-treated populations of MCF7 cancer cells, but not non-cancer cells

The peak wavelength of the LED light used in this study was characterized as 734 nm

(Figure 2A), with a full-width half maxima of 43 nm. The full daily exposure led to a total fluence of 63 mJ/cm^2^, which did not induce significant thermal effects (Figure 2B).

**Figure 2:**
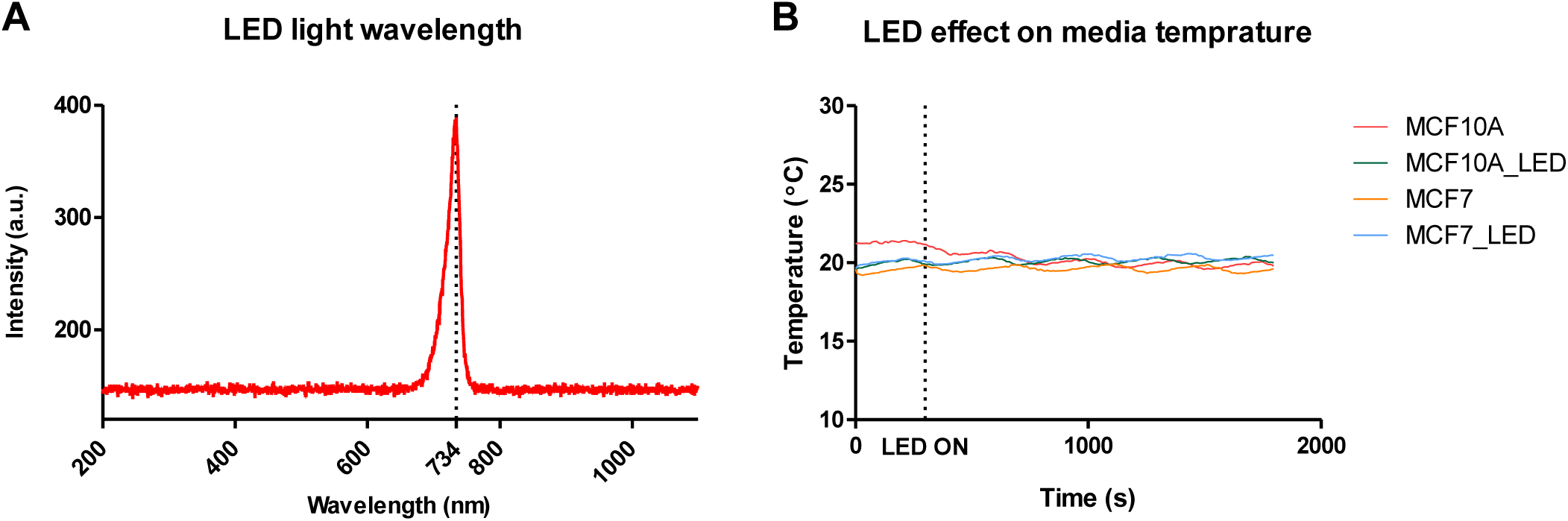
LED light characterisation and effect on mammalian cells (A) The wavelength of LED light count. Peak was identified at 734 nm. Where a.u.; arbitrary units. (B) Temperature measurement during irradiation showing no thermal effect of LED light on cancer and non-cancer cellular media. The LED light was switched on at 300 seconds.

NIR-light exposure of 63 mJ/cm^2^ over a 20-minute period increased senescence in breast cancer cells by 13.5% in Dox-untreated, and 13.7% in Dox-treated populations (P=0.008). However, there was no change in senescence in either non-cancer MCF10A (P=0.97) or IMR-90 (P=0.07) cell lines, in the presence or absence of Dox. Furthermore, in all three cell lines, no Dox – NIR-light significant interaction was observed (Interaction; MCF7: P=0.65, MCF10A: P=0.92, IMR-90: P=0.54; Figure 3).

**Figure 3:**
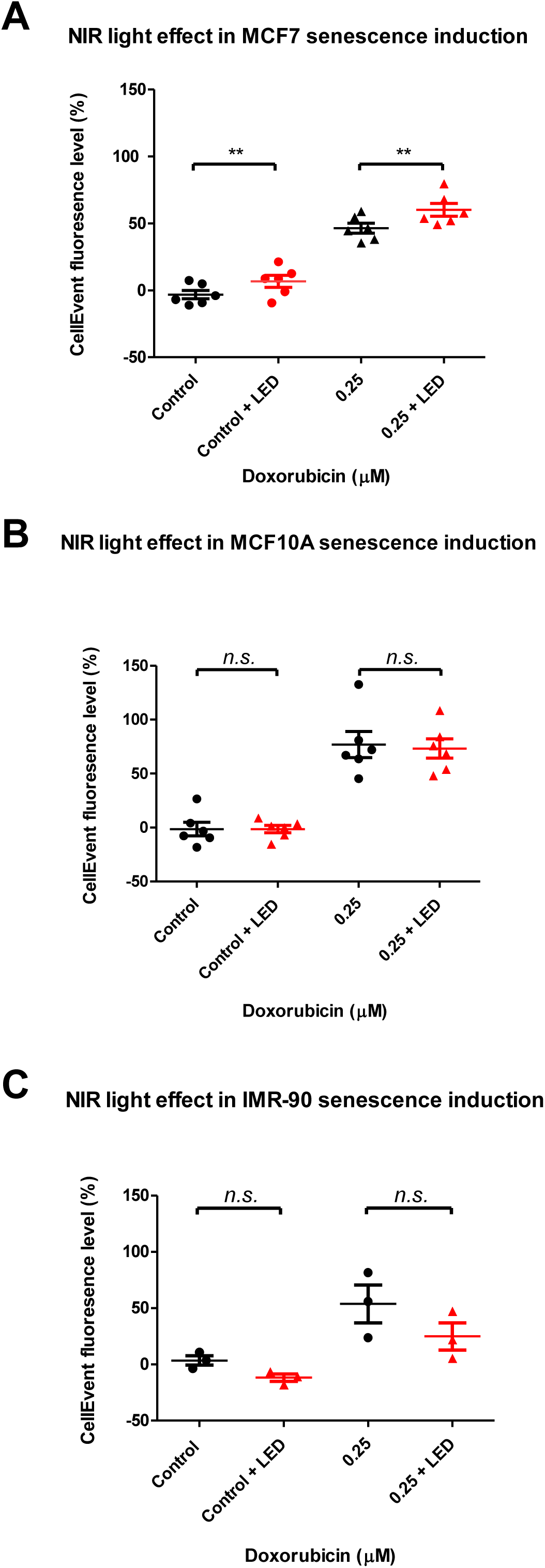
Effect of NIR-light exposure on non-treated and 25 μM Dox-treated cells measured by flow cytometry (A) MCF7 (N=6), (B) MCF10A (N=6) and (C) IMR-90 (N=3) cells were treated with 0 and 0.25 μM Dox and each population was exposed to either NIR-light or no light. Senescent levels were quantified with CellEvent fluorescence intensity. Data are presented as mean changes compared to an untreated control ± SEM. **: P≤0.01, non-significant (n.s.): P>0.05.

### NIR-light treatment increases ROS in MCF7 cancer cells

ROS levels of MCF7 breast cancer cells were assessed in terms of NIR-light exposure, for both Dox-treated and Dox-untreated cell populations. A significant increase in ROS production (P=0.015) was observed after NIR-light treatment, with a 36% increase in Dox-untreated and 94.6% in Dox-treated cells (Figure 4). No Dox – NIR-light significant interaction was observed in terms of ROS levels (P= 0.25).

**Figure 4:**
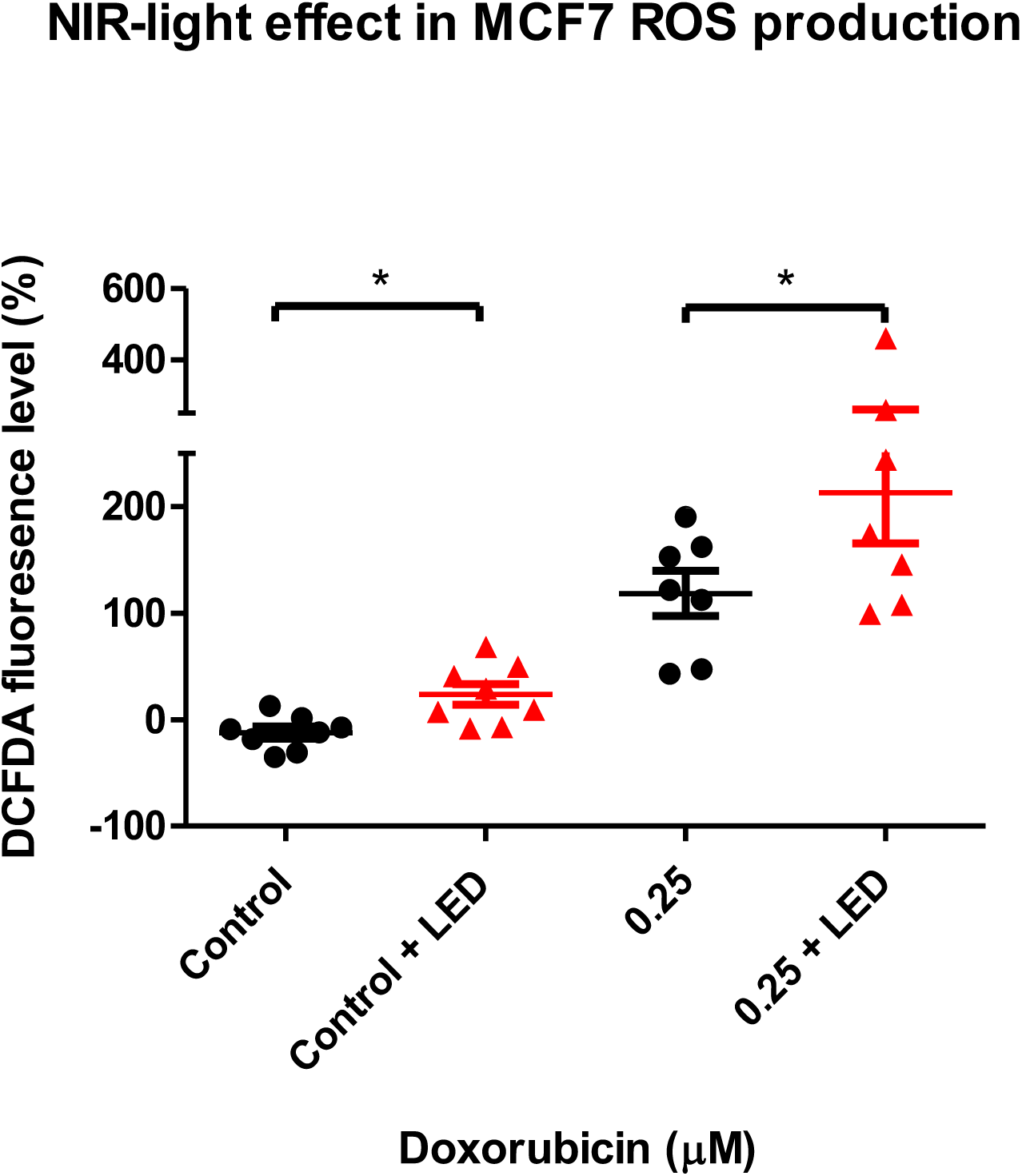
The effect of NIR-light on ROS production measured by flow cytometry MCF7 (N=7) cells were treated with 0 and 0.25 μM Dox and each population was exposed to no and NIR-light. ROS levels were quantified with DCFDA fluorescence intensity. Data are presented as mean changes compared to an untreated control ± SEM. *: P≤ 0.05.

### NIR-light treatment does not affect mitochondrial Ca^2+^ levels in MCF7 cells

Using Rhod2 fluorescence dye, mitochondrial Ca^2+^ levels of MCF7 breast cancer cells were assessed following chronic NIR-light treatment. No significant changes in Ca^2+^ levels were observed in Dox-treated and Dox-untreated populations after NIR exposure (P=0.60; Figure 5). No Dox – NIR-light significant interaction was observed in terms of Ca^2+^ levels (P= 0.84).

**Figure 5:**
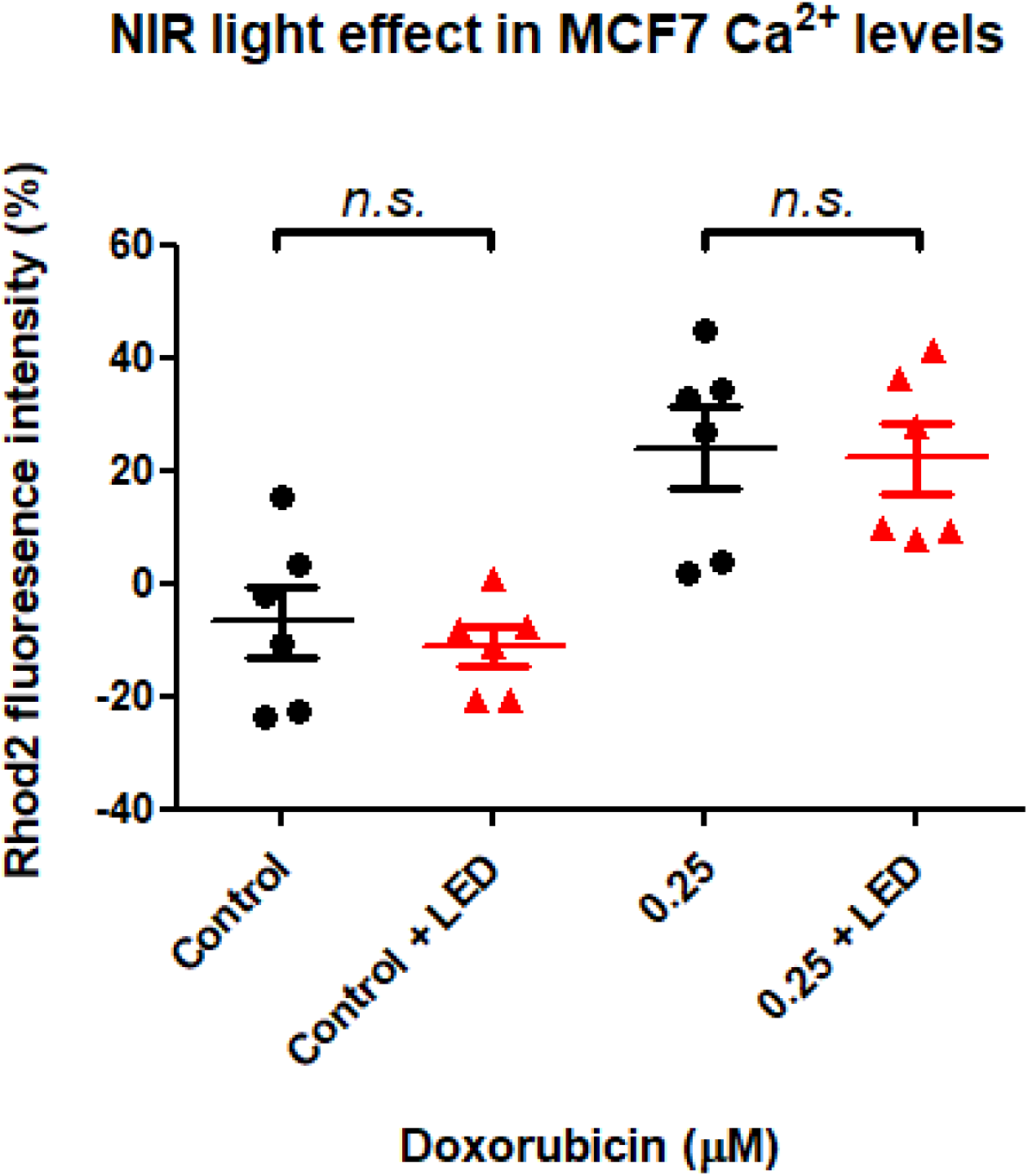
The effect of NIR-light on Ca^2+^ levels measured by flow cytometry MCF7 cells (N=6) were treated with 0 and 0.25 μM Dox and each population was exposed to no and NIR-light. Ca^2+^ levels were quantified with Rhod2 fluorescence intensity. Data are presented as mean changes compared to an untreated control ± SEM. non-significant (n.s.): P>0.05.

### NIR-light treatment increases mitochondrial membrane potential in MCF7 cells

Using TMRE dye, MCF7 breast cancer cells showed a significant increase (P=0.02) in mitochondrial membrane potential (MMP) after chronic NIR-light treatment, 14.9% in Dox-untreated and 5.4% in Dox-treated populations (Figure 6). No Dox – NIR-light significant interaction was observed in terms of MMP levels (P= 0.24).

**Figure 6:**
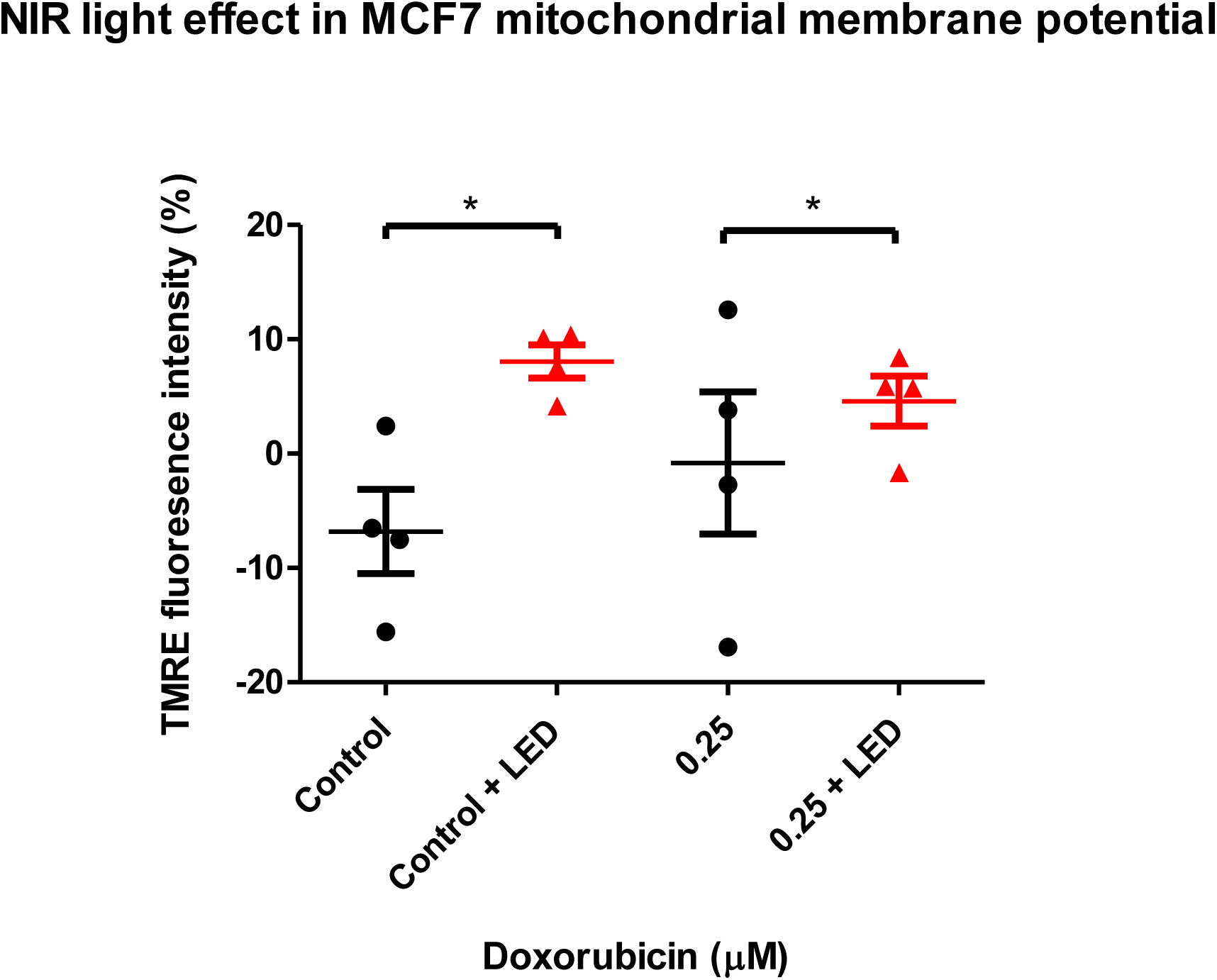
The effect of NIR-light on MMP measured by flow cytometry MCF7 cells (N=4) were treated with 0 and 0.25 μM Dox and each population was exposed to no and NIR-light. MMP was quantified with TMRE fluorescence intensity. Data are presented as mean changes compared to an untreated control ± SEM. *: P≤ 0.05.

## DISCUSSION

### Low intensity NIR-light can induce senescence in MCF7 cancer cells

In this study, the effects of chronic NIR-light treatment on senescence were investigated in three different cell lines (MCF7, MCF10A and IMR-90). Exposure to 734 nm NIR-light, at an intensity of 63 mJ/cm^2^ per day for six days, induced senescence in MCF7 cancer cells, but not in MCF10A or IMR-90 non-cancer cell lines. Moreover, we observed that NIR-light exposure was associated with significant changes in MMP and cellular ROS, in the absence of changes in mitochondrial Ca^2+^ levels.

Previous studies have evaluated the effects of light exposure on cancer cell growth. In these studies a wavelength circa 633 nm was used as light at this wavelength is thought to modulate CCO (Pastore et al., 2000). Melanoma cells (A2058) were shown to increase their growth rate, in a dose-response manner, following exposure to light at 500, 1,000, and 2,000 mJ/cm^2^, with a similar dose-response increase in CCO activity (Hu et al., 2007). Similarly, PBM exposure to 180 mJ/cm^2^ fluence has been reported to increase cell growth in mouse epithelial breast (EMT-6) and mouse sarcoma cell lines (RIF-1) (Al-Watban and Andres, 2012). Furthermore, there are suggestions that the response may be dose-related. For instance, MCF7 cells exposed to 1000 mJ/cm^2^ increased cell number and metabolic activity, but the effect stopped at fluences above 5,000 mJ/cm^2^, while, at a very low fluence, similar to our study (28 mJ/cm^2^), there was no increase in cell proliferation, while elevated metabolic activity remained (Magrini et al., 2012). Though Magrini et al did not assess senescence, the overall phenotype following low doses of light exposure is suggestive of senescence, despite the wavelength used being nominally below the infrared region. This is probably because one of the potential photonic receptors, CCO, has a broad range of absorbances from red to infrared (Golovynska et al., 2022).

Thus, it appears that with exposure to doses of red/NIR-light ∼100 mJ/cm^2^ there is a cessation in proliferation in cancer cells, associated with permanent cell cycle arrest and increased metabolic activity. However, when the fluence of light is increased above this range, there is increased cellular proliferation together with elevated metabolism in cancer cells (Hamblin et al, 2018).

### NIR-light does not affect senescence induction levels in non-cancer cell lines

While cancer cells exhibited a significant permanent cell cycle arrest after NIR-light exposure, there were no comparable changes in non-cancer breast control and lung fibroblast cell lines. This suggests that the mechanism of senescence induction may be cell-dependent (Gorgoulis et al., 2019) and may relate to the inherent metabolic activity of cancer cells, which are known to be more metabolically active than normal cells (Perillo et al., 2020).

Numerous studies have shown that exposure to red/NIR-light at doses >100 mJ/cm^2^ can significantly increase the proliferation of a wide variety of human and non-human non-cancer cell lines (Abrahamse et al., 2010; Kim et al., 2019; Gagnon et al., 2016). In our study we did not observe such increase in cell proliferation in non-cancer cells, probably because the overall exposure dose was significantly below 100 mJ/cm^2^, suggesting that there may be a Goldilocks range at which red/NIR light may induce senescence in cancer cell, but not increase proliferation in non-cancer cell. These results would certainly support earlier studies that indicate that cellular responses to light may be specific to tissue origin and metabolic activity (Laakso et al., 1993).

The potential senescence disparity response between non-cancer and cancer cells, suggests a possible selective effect involving CCO. As previously discussed, CCO appears to serve as a primary chromophore in PBM therapy at red/NIR wavelengths (Hamblin, 2017). Several studies have shown reduced CCO activity in cancer (Srinivasan et al., 2016). Defects in CCO can cause a metabolic shift to glycolysis, and the loss of CCO activity results in the activation of oncogenic signalling pathways and the upregulation of the genes involved in cell signalling, cell migration, and extracellular matrix interactions (Warburg, 1956; Srinivasan et al., 2016). This suggest that cancer and non-cancer cells may respond differently to PBM. Interestingly, in a study using both cancerous and non-cancerous cell lines, the former demonstrated a greater shift towards glycolysis and higher ROS levels after almost three minutes of exposure to 780 nm light even at very low fluence (5 J/cm^2^), which may be explained by the Warburg effect (Gonçalves de Faria et al., 2021; Warburg, 1956). Additionally, 24 hours after treatment, non-cancer cells increased proliferation, whereas cancer cells did not (Gonçalves de Faria et al., 2021). This suggests that at low fluence the initial ROS increase may play a different role in cancer and non-cancer cells, with the latter allowing low ROS levels to modulate pro-survival and protective mechanisms, to overcome stress and escape senescence induction (Gonçalves de Faria et al., 2021).

### NIR-light treatment induces oxidative stress in MCF7 cancer cells

As previously shown, increase in ROS levels is one of the primary effects of PMB (Amaroli et al., 2022). We have confirmed this in our study and have gone further to show that it is closely associated to senescence induction by NIR-light treatment. Moreover, as ROS is one of the hallmarks of cellular senescence, of the induction of senescence by red/NIR light resulted in an even higher ROS increase, probably due to complications of cellular OXPHOS (Höhn et al., 2017).

Mitochondria-nuclei mediators, such as ROS, are known to stimulate the activation of various transcription factors, such as nuclear factor kappa B (NF-kB), or nuclear factor erythroid 2–related factor 2 (Nrf2), which can control the expression of numerous genes responsible for cellular functions, including survival and adaptation, balancing inflammation and anti-oxidant pathways, all of which are key in modulating mitochondrial function (Wang et al., 2002; Brigelius-Flohe and Flohe, 2011). The effect of cellular senescence is activated and promoted by NF-kB activation (Rovillain et al., 2011), while reduced Nrf2 expression is associated with senescence induction (Yuan et al., 2020). Additionally, it has been shown that a burst of ROS due to PBM treatment can activate NF-kB signalling pathway (Hamblin, 2017), both in cancer (Margini et al, 2012) and non-cancer cellular models (Brigelius-Flohe and Flohe, 2011), while, more recent data has shown that PBM modulates Nrf2 in skin immunomodulation (Salman et al., 2023), supporting the concept that PBM is modulating cellular redox and associated transcriptional landscape. Further work is needed to elucidate the exact role of ROS in the modulation of the NF-kB/Nrf2 in cancer cells following red/NIR exposure.

### NIR-light modulates MCF7 cancer mitochondrial membrane potential, but not Ca^2+^ levels

It is well established that mitochondria can modulate nuclear function via retrograde signalling, involving moieties such as ROS, NO and Ca^2+^, which are associated with MMP. At low doses, PBM has been reported to increase their levels (Karu 2008). However, in the present study, only ROS and MMP but not mitochondrial Ca^2+^ levels were influenced by PBM. This confirms the complexity of PBM effects on molecular and cellular mechanistic level (Kujawa et al., 2014); together with the effectiveness of PBM, which depends on several factors, the light treatment parameters and the origin of the tissue/cell line influence differently the molecular mechanisms that alter the transcription factors.

In the present study, MMP was increased in MCF7 cancer Dox-untreated and senescent populations after NIR-light treatment. MMP is a key indicator of mitochondrial status and is constantly adapting to cellular needs, as the energy extracted from metabolism, and is very tightly regulated, hence the close relationship between ROS, Ca^2+^ and uncoupling (Cortassa et al., 2014). Several studies using mitochondria isolated from different models have demonstrated that an increase in ROS production is highly correlated with an increase in MMP (Korshunov et al., 1997; Suski et al., 2012). More recent data shows that red/NIR light exposure stimulates MMP levels and the production of ROS, and the effect depends on the doses. For example, light at 650 and 808 nm between 3 and 30 J/cm^2^ increased the viability of a number of cell lines by 5-10%, whereas above 90 J/cm^2^ exposure decreases cell viability, and increases cell death (Golovynska et al., 2023). Interestingly, although cells after red/NIR light exposure show a biphasic response, they gradually increase their MMP as the fluences increase from 3 to 90 J/cm^2^ (Golovynska et al., 2023). Our data clearly showed that exposure of MCF7 cells to light at 734 nm, at a daily dose of 63.6 mJ/cm^-2^ resulted also in an increase in MMP.

Additionally, decreasing MMP is associated with mitochondrial dysfunction and ROS production considered damaging to the tissue (Hamblin, 2017). This suggests that senescence induction in cancer cell after NIR-light exposure is the result of ROS, as a secondary messenger, and not a general mitochondrial dysfunction.

Although altered MMP was observed in senescent cells in this and other studies (Vasileiou et al., 2019), no change in mitochondrial Ca^2+^ levels were recorded after NIR-light exposure. Mitochondrial Ca^2+^ uptake is known to follow a biphasic effect: uptake can initially stimulate mitochondrial function, resulting in increased ROS and increased membrane potential, and is a normal part of Ca^2+^ signalling. However, too much Ca^2+^ can result in overload and the collapse of the MMP and potentially, cell death. This process is tightly controlled (Rossi et al, 2019). Thus, it is possible that a low dose of light may stimulate mitochondrial ROS, and possibly induce a transient change in mitochondrial Ca^2+^ flux in which case it would not be detectable by the techniques employed in the current study. However, it has been reported that cancer does redirects Ca^2+^ away from mitochondria (Monteith et al., 2017), and that mitochondria exhibit tight control of Ca^2+^ flux to prevent excessive overload. This effect is tightly integrated with the membrane potential, for instance, increased MMP can enhance NADPH production to prevent oxidative stress, while enhancing Ca^2+^ release (Vercesi et al., 2018).

## Conclusion

Our study demonstrates that a selective effect of low-dose PBM at NIR wavelength induces senescence in a cancer cell line without affecting non-cancer cell lines. The mechanism of senescence induction appears to involve disruption of mitochondrial homeostasis, leading to an increase in MMP and subsequent production of ROS, which may act as secondary messengers to interact with the nucleus and promote senescence. Further studies examining NIR-light cell exposure are needed to test the potential implication of the findings reported here, together with studies confirming the signalling pathways involved in this proposed mechanism.

## EXPERIMENTAL PROCEDURES

### Cell Lines

Human breast cell line *MCF10A* was grown in DMEM: F12 (Life Sciences, UK), supplemented with 5% Horse Serum (ThermoFisher, UK), 2% Penicillin/Streptomycin, 20 ng/mL epidermal growth factor (Merck, UK), 0.5 mg/mL hydrocortisone (Merck, UK), 100 ng/mL cholera toxin (Merck, UK) and 10 μg/mL insulin (Merck, UK). Human breast adenocarcinoma cell line *MCF7* and human lung fibroblast cell line *IMR-90* purchased from American Type Culture Collection (ATCC) were grown in MEME (ThermoFisher, UK), supplemented with 10% FBS, 1% l-glutamine and 1% Penicillin/Streptomycin.

### Treatment Protocols for Senescence Induction

Cellular premature senescence was induced by treatment with 0.25 μM Dox (Tocris, Bio-techne, Bristol) for 24 hours and subsequent extended culture in drug-free media for 5 days (Hernandez-Segura et al., 2018a).

### LED NIR-light characterization

LED NIR-light exposure was achieved by using a pre-mounted 7 – LED array of NIR-light (734 nm with a full-width half maxima of 43 nm) Rebel LEDs (Luxeon Star LED, Alberta – Canada) mounted on a SinkPAD-II 40 mm Round 7-Up base with a total power output of between 0.328 to 0.420 mW (power density at sample measured ≈ 0.053 mW/cm^2^, total fluence over 20 minutes was ≈ 63 mJ/cm^2^), as described by (Pooam et al., 2021).

LED light characterization was performed at the Central Laser Facility, Research Complex at Harwell, Science & Technology Facilities Council, UK. The spectrum was measured using a portable USB high-resolution fiber optic HR2000CG UV-NIR spectrometer (Ocean Optics Inc, Florida – USA), and the output power was recorded on a calibrated PM100D power meter with an S120VC photodiode power sensor (Thorlabs Inc, New Jersey – USA). The thermal change of the media exposed to LED light was measured with a FireSting optical temperature sensor, recorded by an FSO2 temperature meter (Pyroscience, Germany).

### NIR-light light treatment

The cells were seeded in two 6-well plates. One plate (“control”) was left in the dark and a second (“treated”) was exposed to the NIR-light at about 0.053 mW/cm^2^ for 20 minutes per day, for the next 6 days, which gave a daily dose of 63 mJ/cm^2^ and a total dose of 381.6 mJ/cm^2^. The LED array was placed 40 mm below the 6-well culture plate and in between 2 wells, to create a uniform beam for illumination of the culture plate. The well with the control “untreated” population was covered and any other cell treatment (eg. Dox treatment) was performed normally in the period of these 6 days, before the light treatment.

### Flow cytometry assay for senescence quantification

Flow cytometry was used for fluorescent quantification and for examining senescent levels in cellular populations (Cahu and Sola, 2013). CellEvent™ Senescence Green Flow Cytometry Assay Kit (ThermoFisher, UK) was used for flow cytometric detection of cellular senescence via using a fluorescent dye that binds to the senescence-associated beta-galactosidase (SA-beta-gal) enzyme, and the assay was performed according to manufacture protocol (ThermoFisher Scientific, 2019).

Flow cytometry measurements were performed with a BD LSRFortessa^TM^ X-20 Analyser flow cytometer (BD Biosciences, New Jersey-USA) and data were recorded/analyzed using a BD FACSDiva™ Software. The AlexaFluor channel (488 nm laser) was used to capture the uptake of stained cells. For the analysis, the cell population was selected on a forward/scatter/sideward scatter plot to exclude debris cellular aggregates, and 10,000 gated events were recorded. The median intensity was determined from a histogram and results were recorded as a median value.

### Detection and Quantification of ROS

The 2’,7’ –dichlorofluorescin diacetate (DCFDA) stain was used to detect changes in the levels of ROS in live cells. After treatment cells were washed with PBS and incubated with 30 µM DCFDA (Sigma, UK) at 37°C 5% CO_2_ for 30 minutes, protected from light. Fluorescence was quantified by flow cytometry –the AlexaFluor channel (488-nm laser) was used to capture the uptake of stained cells.

### Assessment of cellular Ca^2+^ levels

Rhod-2 (ThermoFisher, UK) was used to label mitochondrial Ca^2+^ in live cells. After treatment, cells were incubated with 1 μM Rhod-2 in Phenol Red Free media, at 37°C 5% CO_2_ for 30 minutes, protected from light. Rhod-2 fluorescence was quantified by flow cytometry – the PE channel (575 nm laser) was used to capture the uptake of stained cells.

### Assessment of MMP

The cell-permeant tetramethylrhodamine, ethyl ester (TMRE) stain was used to label active mitochondria in live cells. After treatment cells were incubated with 500 nM TMRE (Sigma, UK) in Phenol Red Free media, at 37°C 5% CO_2_ for 30 minutes, protected from light. Active mitochondria sequester TMRE and were quantified by flow cytometry – the PE channel (575 nm laser) was used to capture the uptake of stained cells.

### Statistical Analysis

Results are expressed as the mean ± standard error of the mean (SEM), at a minimum of 3 biological repeats. Where appropriate, outliers were detected and excluded using Grubb’s Test.

For comparison between two groups, a two-tailed, unpaired T-test (F-test; P-value) was applied. The effect of NIR-light treatment in replicative control and senescent populations in senescence induction; ROS levels; and Ca^2+^ levels were compared with a mixed-model two-way ANOVA, with a Bonferonni *post hoc* test.

## Author contribution

IK, RM, SW, AL conceived and carried out the experiments. IK, RM analyzed the data. IK, RM, AN, JB contributed on conceptualization. All authors were involved in writing and had final approval of the manuscript.

## Data Availability Statement

Data available on request from the authors.

## Bibliography

1. Abrahamse, H., Houreld, N.N., Muller, S., Ndlovu, L., 2010. Fluence and wavelength of low intensity laser irradiation affect activity and proliferation of human adipose derived stem cells. Medical Technology SA 24.

2. Al-Watban, F.A.H., Andres, B.L., 2012. Laser biomodulation of normal and neoplastic cells. Lasers Med Sci 27, 1039–1043. https://doi.org/10.1007/s10103-011-1040-9

3. Amaroli, A., Pasquale, C., Zekiy, A., Benedicenti, S., Marchegiani, A., Sabbieti, M.G., Agas, D., 2022. Steering the multipotent mesenchymal cells towards an anti-inflammatory and osteogenic bias via photobiomodulation therapy: How to kill two birds with one stone. J Tissue Eng 13, 204173142211101. https://doi.org/10.1177/20417314221110192

4. Ash, C., Dubec, M., Donne, K., Bashford, T., 2017. Effect of wavelength and beam width on penetration in light-tissue interaction using computational methods. Lasers Med Sci 32, 1909–1918. https://doi.org/10.1007/s10103-017-2317-4

5. Begum, R., Powner, M.B., Hudson, N., Hogg, C., Jeffery, G., 2013. Treatment with 670 nm Light Up Regulates Cytochrome C Oxidase Expression and Reduces Inflammation in an Age-Related Macular Degeneration Model. PLoS One 8, e57828. https://doi.org/10.1371/journal.pone.0057828

6. Brigelius-Flohé, R., Flohé, L., 2011. Basic Principles and Emerging Concepts in the Redox Control of Transcription Factors. Antioxidants & Redox Signaling, 15(8), 2335–2381. https://doi.org/10.1089/ars.2010.3534

7. Chen, A.C.-H., Arany, P.R., Huang, Y.-Y., Tomkinson, E.M., Sharma, S.K., Kharkwal, G.B., Saleem, T., Mooney, D., Yull, F.E., Blackwell, T.S., Hamblin, M.R., 2011. Low-Level Laser Therapy Activates NF-kB via Generation of Reactive Oxygen Species in Mouse Embryonic Fibroblasts. PLoS One 6, e22453. https://doi.org/10.1371/journal.pone.0022453

8. Choi, M., Na, S.Y., Cho, S., Lee, J.H., 2011. Low Level Light Could Work on Skin Inflammatory Disease: A Case Report on Refractory Acrodermatitis Continua. J Korean Med Sci 26, 454. https://doi.org/10.3346/jkms.2011.26.3.454

9. Choi, W., Baik, K.Y., Jeong, S., Park, S., Kim, J.E., Kim, H.B., Chung, J.H., 2021. Photobiomodulation as an antioxidant substitute in post-thawing trauma of human stem cells from the apical papilla. Sci Rep 11, 17329. https://doi.org/10.1038/s41598-021-96841-3

10. Cortassa, S., Sollott, S. J., & Aon, M. A., 2017. Mitochondrial respiration and ROS emission during β-oxidation in the heart: An experimental-computational study. PLOS Computational Biology, 13(6), e1005588. https://doi.org/10.1371/journal.pcbi.1005588

11. da Silva, P.F.L., Ogrodnik, M., Kucheryavenko, O., Glibert, J., Miwa, S., Cameron, K., Ishaq, A., Saretzki, G., Nagaraja-Grellscheid, S., Nelson, G., von Zglinicki, T., 2019. The bystander effect contributes to the accumulation of senescent cells in vivo. Aging Cell 18, e12848. https://doi.org/10.1111/acel.12848

12. de Sousa, M. V. P., 2017. What is low-level laser (light) therapy?, in: Michael R. Hamblin, Tanupriya Agrawal, Marcelo de Sousa (Eds.), Handbook of Low-Level Laser Therapy. Pan Stanford Publishing Pte. Ttd., Sao Paolo.

13. Gagnon, D., Gibson, T. W. G., Singh, A., zur Linden, A. R., Kazienko, J. E., & LaMarre, J. (2016). An in vitro method to test the safety and efficacy of low-level laser therapy (LLLT) in the healing of a canine skin model. BMC Veterinary Research, 12(1), 73. https://doi.org/10.1186/s12917-016-0689-5

14. Golovynska, I., Golovynskyi, S., & Qu, J., 2023. Comparing the Impact of <scp>NIR</scp>, Visible and <scp>UV</scp> Light on <scp>ROS <scp> Upregulation via Photoacceptors of Mitochondrial Complexes in Normal, Immune and Cancer Cells. Photochemistry and Photobiology, 99(1), 106–119. https://doi.org/10.1111/php.13661

15. Gonçalves de Faria, C.M., Ciol, H., Salvador Bagnato, V., Pratavieira, S., 2021. Effects of photobiomodulation on the redox state of healthy and cancer cells. Biomed Opt Express 12, 3902. https://doi.org/10.1364/BOE.421302

16. Gorgoulis, V., Adams, P.D., Alimonti, A., Bennett, D.C., Bischof, O., Bishop, C., Campisi, J., Collado, M., Evangelou, K., Ferbeyre, G., Gil, J., Hara, E., Krizhanovsky, V., Jurk, D., Maier, A.B., Narita, M., Niedernhofer, L., Passos, J.F., Robbins, P.D., Schmitt, C.A., Sedivy, J., Vougas, K., von Zglinicki, T., Zhou, D., Serrano, M., Demaria, M., 2019. Cellular Senescence: Defining a Path Forward. Cell 179, 813–827. https://doi.org/10.1016/j.cell.2019.10.005

17. Hamblin, M.R., 2017. Mechanisms and Mitochondrial Redox Signaling in Photobiomodulation. Photochem Photobiol 94, 199–212. https://doi.org/10.1111/php.12864

18. Hamblin, M.R., Nelson, S.T., Strahan, J.R., 2018. Photobiomodulation and Cancer: What Is the Truth? Photomed Laser Surg 36, 241–245. https://doi.org/10.1089/pho.2017.4401

19. Hayflick, L., Moorhead, P.S., 1961. The serial cultivation of human diploid cell strains. Exp Cell Res 25, 585–621. https://doi.org/10.1016/0014-4827(61)90192-6

20. He, S., Sharpless, N.E., 2017. Senescence in Health and Disease. Cell 169, 1000–1011. https://doi.org/10.1016/j.cell.2017.05.015

21. Höhn, A., Weber, D., Jung, T., Ott, C., Hugo, M., Kochlik, B., Kehm, R., König, J., Grune, T., Castro, J.P., 2017. Happily (n)ever after: Aging in the context of oxidative stress, proteostasis loss and cellular senescence. Redox Biol 11, 482–501. https://doi.org/10.1016/j.redox.2016.12.001

22. Hu, W.-P., Wang, J.-J., Yu, C.-L., Lan, C.-C.E., Chen, G.-S., Yu, H.-S., 2007. Helium–Neon Laser Irradiation Stimulates Cell Proliferation through Photostimulatory Effects in Mitochondria. Journal of Investigative Dermatology 127, 2048–2057. https://doi.org/10.1038/sj.jid.5700826

23. Karu, T.I., 2008. Mitochondrial Signaling in Mammalian Cells Activated by Red and Near-IR Radiation. Photochem Photobiol 84, 1091–1099. https://doi.org/10.1111/j.1751-1097.2008.00394.x

24. Kim, K., Lee, J., Jang, H., Park, S., Na, J., Myung, J., Kim, M.-J., Jang, W.-S., Lee, S.-J., Kim, H., Myung, H., Kang, J., Shim, S., 2019. Photobiomodulation Enhances the Angiogenic Effect of Mesenchymal Stem Cells to Mitigate Radiation-Induced Enteropathy. Int J Mol Sci 20, 1131. https://doi.org/10.3390/ijms20051131

25. Korshunov, S.S., Skulachev, V.P., Starkov, A.A., 1997. High protonic potential actuates a mechanism of production of reactive oxygen species in mitochondria. FEBS Lett 416, 15–18. https://doi.org/10.1016/S0014-5793(97)01159-9

26. Kujawa, J., Pasternak, K., Zavodnik, I., Irzmański, R., Wróbel, D., Bryszewska, M., 2014. The effect of near-infrared MLS laser radiation on cell membrane structure and radical generation. Lasers Med Sci 29, 1663–1668. https://doi.org/10.1007/s10103-014-1571-y

27. Laakso, L., Richardson, C., Cramond, T., 1993. Factors affecting Low Level Laser Therapy. Australian Journal of Physiotherapy 39, 95–99. https://doi.org/10.1016/S0004-9514(14)60473-6

28. Magrini, T.D., dos Santos, N.V., Milazzotto, M.P., Cerchiaro, G., da Silva Martinho, H., 2012. Low-level laser therapy on MCF-7 cells: a micro-Fourier transform infrared spectroscopy study. J Biomed Opt 17, 1015161. https://doi.org/10.1117/1.JBO.17.10.101516

29. Mester, E., Ludany, G., Selyei, M., Szende, B., Total, G.J., 1968. The stimulating effect of low power laser rays on biological systems. Laser Rev. 1.

30. McHugh, D., Gil, J., 2018. Senescence and aging: Causes, consequences, and therapeutic avenues. Journal of Cell Biology 217, 65–77. https://doi.org/10.1083/jcb.201708092

31. Monteith, G. R., Prevarskaya, N., & Roberts-Thomson, S. J., 2017. The calcium–cancer signalling nexus. Nature Reviews Cancer, 17(6), 373–380. https://doi.org/10.1038/nrc.2017.18

32. Pastore, M., Greco, S., Passarella, D., 2000. Specific helium-neon laser sensitivity of the purified cytochrome c oxidase. International Journal of Radiation Biology, 76(6), 863– 870. https://doi.org/10.1080/09553000050029020

33. Perillo, B., Di Donato, M., Pezone, A., Di Zazzo, E., Giovannelli, P., Galasso, G., Castoria, G., Migliaccio, A., 2020. ROS in cancer therapy: the bright side of the moon. Exp Mol Med 52, 192–203. https://doi.org/10.1038/s12276-020-0384-2

34. Pooam, M., Aguida, B., Drahy, S., Jourdan, N., Ahmad, M., 2021b. Therapeutic application of light and electromagnetic fields to reduce hyper-inflammation triggered by COVID-19. Commun Integr Biol 14, 66–77. https://doi.org/10.1080/19420889.2021.1911413

35. Robijns, J., Nair, R. G., Lodewijckx, J., Arany, P., Barasch, A., Bjordal, J. M., Bossi, P., Chilles, A., Corby, P. M., Epstein, J. B., Elad, S., Fekrazad, R., Fregnani, E. R., Genot, M.-T., Ibarra, A. M. C., Hamblin, M. R., Heiskanen, V., Hu, K., Klastersky, J., … Bensadoun, R.-J., 2022. Photobiomodulation therapy in management of cancer therapy-induced side effects: WALT position paper 2022. Frontiers in Oncology, 12. https://doi.org/10.3389/fonc.2022.927685

36. Rossi, A., Pizzo, P., & Filadi, R., 2019. Calcium, mitochondria and cell metabolism: A functional triangle in bioenergetics. Biochimica et Biophysica Acta (BBA) – Molecular Cell Research, 1866(7), 1068–1078. https://doi.org/10.1016/j.bbamcr.2018.10.016

37. Rovillain, E., Mansfield, L., Caetano, C., Alvarez-Fernandez, M., Caballero, O.L., Medema, R.H., Hummerich, H., Jat, P.S., 2011. Activation of nuclear factor-kappa B signalling promotes cellular senescence. Oncogene 30, 2356–2366. https://doi.org/10.1038/onc.2010.611

38. Salman, S., Guermonprez, C., Peno-Mazzarino, L., Lati, E., Rousseaud, A., Declercq, L.,; Kerdine-Römer, S., 2023. Photobiomodulation Controls Keratinocytes Inflammatory Response through Nrf2 and Reduces Langerhans Cells Activation. Antioxidants, 12(3), 766. https://doi.org/10.3390/antiox12030766

39. Sommer, A.P., Haddad, M.Kh., Fecht, H.-J., 2015. Light Effect on Water Viscosity: Implication for ATP Biosynthesis. Sci Rep 5, 12029. https://doi.org/10.1038/srep12029

40. Srinivasan, S., Guha, M., Dong, D. W., Whelan, K. A., Ruthel, G., Uchikado, Y., Natsugoe, S., Nakagawa, H., & Avadhani, N. G. (2016). Disruption of cytochrome c oxidase function induces the Warburg effect and metabolic reprogramming. Oncogene, 35(12), 1585– 1595. https://doi.org/10.1038/onc.2015.227

41. Suski, J.M., Lebiedzinska, M., Bonora, M., Pinton, P., Duszynski, J., Wieckowski, M.R., 2012. Relation Between Mitochondrial Membrane Potential and ROS Formation. pp. 183–205. https://doi.org/10.1007/978-1-61779-382-0_12

42. Vasileiou, P., Evangelou, K., Vlasis, K., Fildisis, G., Panayiotidis, M., Chronopoulos, E., Passias, P.-G., Kouloukoussa, M., Gorgoulis, V., Havaki, S., 2019. Mitochondrial Homeostasis and Cellular Senescence. Cells 8, 686. https://doi.org/10.3390/cells8070686

43. Vercesi, A. E., Castilho, R. F., Kowaltowski, A. J., de Oliveira, H. C. F., de Souza-Pinto, N. C., Figueira, T. R., & Busanello, E. N. B., 2018. Mitochondrial calcium transport and the redox nature of the calcium-induced membrane permeability transition. Free Radical Biology and Medicine, 129, 1–24. https://doi.org/10.1016/j.freeradbiomed.2018.08.034

44. Wang, T., Zhang, X., Li, J.J., 2002. The role of NF-κB in the regulation of cell stress responses. Int Immunopharmacol 2, 1509–1520. https://doi.org/10.1016/S1567-5769(02)00058-9

45. Warburg, O., 1956. On Respiratory Impairment in Cancer Cells. Science (1979) 124, 269–270. https://doi.org/10.1126/science.124.3215.269

46. Yuan, H., Xu, Y., Luo, Y., Wang, N.-X., & Xiao, J.-H. (2021). Role of Nrf2 in cell senescence regulation. Molecular and Cellular Biochemistry, 476(1), 247–259. https://doi.org/10.1007/s11010-020-03901-9

